# MAPK-mediated PHGDH induction is essential for melanoma formation and represents an actionable vulnerability

**DOI:** 10.1101/2024.04.11.589139

**Authors:** Neel Jasani, Xiaonan Xu, Benjamin Posorske, Yumi Kim, Olga Vera, Kenneth Y. Tsai, Gina M. DeNicola, Florian A. Karreth

## Abstract

Overexpression of PHGDH, the rate-limiting enzyme in the serine synthesis pathway, promotes melanomagenesis, melanoma cell proliferation, and survival of metastases in serine-low environments such as the brain. While *PHGDH* amplification explains PHGDH overexpression in a subset of melanomas, we find that PHGDH levels are universally increased in melanoma cells due to oncogenic BRAF^V600E^ promoting *PHGDH* transcription through mTORC1-mediated translation of ATF4. Importantly, PHGDH expression was critical for melanomagenesis as depletion of *PHGDH* in genetic mouse models blocked melanoma formation. Despite BRAF^V600E^- mediated upregulation, PHGDH was further induced by exogenous serine restriction. Surprisingly, BRAF^V600E^ inhibition diminished serine restriction-mediated PHGDH expression by preventing ATF4 induction, creating a potential vulnerability whereby melanoma cells could be specifically starved of serine by combining BRAF^V600E^ inhibition with exogenous serine restriction. Indeed, we show that this combination promoted cell death in vitro and attenuated melanoma growth in vivo. This study identified a melanoma cell-specific PHGDH-dependent vulnerability.

## INTRODUCTION

Advances in treatments for malignant melanoma, the most aggressive form of skin cancer, have vastly improved patient outcomes and now include both targeted therapies and immune therapies. However, inherent and required resistance hamper the success of these therapies, rendering malignant melanoma a fatal disease in >40% of cases. Understanding the molecular and genetic underpinnings of melanoma drivers and vulnerabilities is therefore needed to identify actionable targets for improved melanoma therapies. One such potential target is Phosphoglycerate Dehydrogenase (PHGDH), the first and rate-limiting enzyme in the Serine Synthesis Pathway (SSP) that catalyzes glycolytic intermediate 3-phospho glycerate to 3-phosphohydroxypyruvate which is ultimately converted to serine. Serine generated de novo via the SSP serves as a precursor for various biomolecules like proteins, nucleotides, and lipids and contributes to the antioxidant defense system via the generation of glutathione (1, 2). Cancer cells exhibit a high demand for serine to support their increased proliferation and the upregulation of SSP enzymes can satisfy this demand.

*PHGDH* is recurrently amplified in melanoma and other cancers in a peak region on chromosome 1p (3, 4). Human melanoma cell lines harboring the 1p amplicon are addicted to PHGDH expression for proliferation (3). An oncogenic role for PHGDH in melanoma has been demonstrated in a Braf^V600E^; Pten^FL/FL^; Tyr-CreERt2 mouse model where PHGDH overexpression resulted in a moderate acceleration of melanomagenesis (5). PHGDH overexpression may also cooperate with Braf^V600E^ in the absence of Pten alterations to promote melanoma formation at very low penetrance (5), while PHGDH overexpression alone was not sufficient to induce melanoma (6). Moreover, endogenous PHGDH is upregulated during melanoma progression (7). This upregulation enables metastasis to the brain, an environment where extracellular Serine is scarce, and sensitizes melanoma brain metastases to PHGDH inactivation. BRAF and NRAS mutant melanoma cells increase PHGDH expression upon developing resistance to the BRAF inhibitor Vemurafenib and the MEK inhibitor PD0325901, respectively (8, 9). Genetic silencing of PHGDH re-sensitizes Vemurafenib-resistant and MEK inhibitor-resistant melanoma cells to suppression of the MAPK pathway (8, 9). Thus, upregulation of PHGDH expression appears to play a critical role during melanoma progression, metastasis, and the development of resistance to targeted therapies. However, whether PHGDH is required for melanomagenesis, and thus a potential therapeutic target, is unknown.

In this study, we show that PHGDH is universally upregulated in melanoma compared to nevi and melanocytes and that PHGDH depletion, but not dietary serine restriction, protects from melanoma formation in vivo. Moreover, we find that BRAF^V600E^ promotes the transcription of *PHGDH* via MAPK/mTORC1 signaling-mediated control of ATF4 translation. BRAF^V600E^ inhibition reduces baseline and serine starvation-induced PHGDH expression to create a melanoma cell-specific vulnerability where BRAF^V600E^ inhibition sensitizes to serine restriction. We demonstrate that the combination of Vemurafenib with extracellular serine depletion promotes cell death in vitro and attenuates melanoma growth in vivo.

## MATERIALS AND METHODS

### TCGA and DepMap

*PHGDH* expression and copy number data from 363 Skin Cutaneous Melanoma (SKCM) samples from The Cancer Genome Atlas (TCGA) were downloaded from cBioportal (cBioportal.org). *PHGDH* expression from RNA sequencing was plotted against copy number variations in Prism. *PHGDH* expression and copy number data for melanoma cell lines were downloaded from DepMap (https://depmap.org/portal/) and correlation analysis was performed. A copy number (relative to ploidy + 1) greater than 1.5 was considered as cutoff for *PHGDH* amplification.

### Cell lines and culture conditions

All human and mouse melanoma cell lines were cultured in RPMI 1640 (Corning, Cat. # 10-040-CV) with 5% FBS (Sigma, Cat. #F0926) at 37°C and 5% CO2. HEK293T Lenti-X cells were cultured in DMEM (Corning, Cat. #10-013-CV) with 10% FBS. All cell lines were authenticated and routinely tested to be mycoplasma-free using the MycoAlert Mycoplasma Detection Kit (Lonza, Cat # LT07-710) every two months. Melanocytes (Hermes1, Hermes2, Hermes3A, and Hermes4B) were cultured in RPMI 1640 containing 10% FBS, 10nM Endothelin-1, 200pM Cholera toxin, 10ng/ml Stem cell factor, and 200nM 12-tetradecanoylphorbol 13-acetateat (TPA) at 37°C and 10% CO2 as described by the Welcome Trust Functional Genomics Cell Bank (https://www.sgul.ac.uk/about/our-institutes/neuroscience-and-cell-biology-research-institute/genomics-cell-ban). Hermes1 expressing BRAF^V600E^ was cultured without TPA. For Ser/Gly depletion experiments, cells were cultured in RPMI 1640 without Serine or Glycine (Thermo Fisher Scientific, Cat. # 50-227-4211), containing 5% dialyzed Fetal Bovine serum (Thermo Fisher Scientific, Cat. # 26400044) and 1% glucose, and Serine (0.285mM) and Glycine (0.13mM) were added back for the control condition.

### Plasmids, lentivirus production and siRNA transfections

shScramble and shPHGDH#1/2 in pLKO.1 targeting human PHGDH were described previously (10). HEK239T Lenti-X cells were transfected with packaging and helper plasmids Δ8.2 and VSVG along with lentiviral vectors at a ratio 9:8:1 using jetPRIME transfection reagent (Polypus, Cat. # 101000046). Lentiviral supernatants were collected 48 hours after transfection followed by centrifugation and filtration through a 0.45 μm syringe filter. Melanoma cells were transduced with lentiviruses in the presence of 8μg/ml polybrene. For siRNA transfections, cells were transfected with 25nM of either non-targeting siRNA (SMARTPool, Horizon Discovery Cat. # D-001810-10-05) or siATF4 (SMARTpool, Horizon Discovery Cat. # L-005125-00-0005) using jetPRIME transfection reagent overnight. The medium was refreshed, and the cells were collected 3 days after transfection for protein and RNA isolation. To generate the HA-ATF4 overexpression construct, we first replaced Puromycin with a Blasticidin resistance marker in LT3GEPIR (Addgene plasmid # 111177), a gift from Johannes Zuber (11), by InFusion cloning. We then replaced the GFP-miR-E cassette with HA-tagged ATF4 cDNA from pLenti-hygro- HA-ATF4 (provided by Lixin Wan) by InFusion cloning. Melanoma cells were stably infected with the lentivirus containing the HA-ATF4 overexpression plasmid and selected with 5µg/ml of Blasticidin for a week. The pLenti-ATF4-uORF-GFP reporter plasmid was generated by placing the ATF4 5’UTR from Addgene plasmid #21850, a gift from David Ron (12), upstream of GFP in pLenti-GFP-Puro (Addgene plasmid # 17448, a gift from Eric Campeau and Paul Kaufman (13)) by InFusion cloning.

### Proliferation assay

Cells were plated at a density of 1,000-3,000 cells/well in 96-well plates 5 days after transducing with pLKO.1 shScramble or shPHGDH#1/2. For the Ser/Gly depletion proliferation assays, cells were plated at equal densities in Ser/Gly-containing or Ser/Gly- deficient medium. The media were refreshed after 3 days. The cells were washed with PBS once and fixed using 4% Paraformaldehyde (VWR, Cat. # 97061-850) for 10 minutes. The cells were then stained with 0.5% crystal violet in 20% methanol. The excess crystal violet was washed gently with distilled water and dried overnight. Crystal violet was extracted using 10% acetic acid and absorbance was measured at 600nM using a Glomax microplate reader.

### Clonogenic Cell Survival Assay

Melanoma cells were plated at a low density of 5,000/well for 1205Lu, SKMel28 and 2,000/well for A375 in 12-well plates in Ser/Gly-containing medium. The next day, cells were treated with DMSO or PLX4032 (0.25μM for 1205Lu, 0.5μM for A375, 1μM SKMel28) in Ser/Gly-containing or Ser/Gly-deficient medium. The media with the drug were refreshed every 3 days. The cells were fixed with 4% Paraformaldehyde once they became confluent after approximately 2 weeks for 1205Lu and 10 days for A375 and SKMel28. The cells were then stained with 0.5% crystal violet, washed with water to remove excess dye, and air dried overnight. The plates were then scanned and analyzed in ImageJ to calculate the area covered by colonies.

### Reporter assay for ATF4 translation

Melanoma cells stably expressing the lentiviral reporter construct were treated with either DMSO, PLX4032 (0.25μM), Temsirolimus (0.1μM), or GCN2iB (20μM) for 48 hours in Ser/Gly-containing or Ser/Gly-deficient medium. After 48 hours, the cells were scraped in PBS, and protein was extracted for western blot analysis. GFP expression was used as a readout for ATF4 translation.

### RT-qPCR

RNA was extracted from cells using TRIzol (Invitrogen, Cat. # 15596018) as per the manufacturer’s protocol. cDNA was generated from 500ng of RNA using PrimeScript RT Master Mix (Takara, Cat. # RR036B). The cDNA was diluted 1:20 and 1µl of the diluted cDNA was used for qPCR analysis. qPCRs were performed using PerfeCTa SYBR green Fastmix (Quantabio, Cat. #101414-276) on a StepOnePlus™ real-time PCR system (Thermo Fisher). 18S or β-actin were used as the internal loading control for qPCR. The relative expression was calculated using the comparative threshold cycle method (2^-1ΔΔct^).

### Droplet digital PCR (ddPCR)

PHGDH (Hs00008098_cn, Thermo Fisher, Cat. # 4400291), or RNAse P (Thermo Fisher, Cat. # 4403326) copy number Taqman probes, ddPCR supermix (Bio-Rad, Cat. # 1863023), buffer, cDNA (100ng/reaction), and water were mixed for a total volume of 22μl per reaction in 96-well plates. The plates were sealed, and droplets were generated using a C1000 Thermocycler under the cycling conditions mentioned in the supplementary datasheet. The plates were then analyzed using a QX-200 (Bio-Rad) Reader to quantify positive and negative droplets using fluorescent signals. Results were quantified using Quantasoft software. The number of positive droplets in a sample determines the concentration of the target in copies/μl of the final reaction.

### Immunoblotting

Cells were washed and scraped in PBS, centrifuged and the pellet was lysed using RIPA buffer containing protease and phosphatase inhibitor cocktail (Thermo Scientific, Cat. # 78440). 25-30µg of protein was used for SDS PAGE. Tumor tissues were dissociated in RIPA buffer with protease and phosphatase inhibitor cocktail using a pellet pestle followed by sonication for 10 minutes. All primary antibodies were diluted in 5% milk in Tris-buffered Saline/0.1% Tween20 and incubated overnight at 4°C. The following primary antibodies were used: PHGDH (Sigma, Cat. #HPA021241), ATF4 (Cell Signaling Technologies, Cat. #11815S), pERK (T202/204) (Cell Signaling Technologies, Cat. # 9101S), ERK (Cell Signaling Technologies, Cat. # 4695S), pS6 Ribosomal protein (S235/236) (Cell Signaling Technologies, Cat. # 4858S), S6 Ribosomal protein (Cell Signaling Technologies, Cat. # 2217S), p4EBP1 (S65) (Cell Signaling Technologies, Cat. # 9451S), 4EBP1 (Cell Signaling Technologies, Cat. # 9452S), HA-Tag (Cell Signaling Technologies, Cat. # 3724S), pEIF2α (S51) (Cell Signaling Technologies, Cat. # 3398), EIF2a (Cell Signaling Technologies, Cat. # 5324), GFP (Cell Signaling Technologies, Cat. # 2956S), BRAF (Sigma, Cat. # HPA001328) and β-actin (Invitrogen, Cat. # AM4302)

### Immunohistochemistry

Tumor tissues were fixed in 10% buffered formalin overnight and dehydrated in 70% ethanol. Tissues were paraffin-embedded, sectioned, and hematoxylin and eosin stained by IDEXX BioAnalytics (Columbia, MO). The tissue sections were de-paraffinized and rehydrated in an alcohol series. Antigen retrieval was performed by heating the sections followed by blocking the endogenous peroxidase with 3% hydrogen peroxide. Immunohistochemistry was performed using VECTASTAIN Elite ABC Kit (Vector Laboratories, Cat. # PK-6100) as per the manufacturer’s instructions and then incubated with DAB peroxidase substrate (Cat. # SK4105). The tissue sections were then counter stained in hematoxylin. Antibodies against Phgdh (1:800, Sigma-Aldrich, HPA021241), pERK (1:300, Cell Signaling Technologies, Cat. # 4370), Cleaved Caspase 3 (1:200, Cell Signaling Technologies, Cat. # 9661) and Ki-67 (1:100, Abcam, Cat. # ab15580) were used for immunohistochemistry.

### Cell Death Analysis

Cell death was analyzed using SYTOX green dye (Thermo Fisher, Cat. # S7020). Cells were plated in 24-well plates at a density of 10,000 (1205Lu) or 5,000 (A375) cells/well. The cells were treated with PLX4032 and SYTOX green dye (20nM) was added to the medium to analyze cell death for 3 days. The plates were placed in a Cellcyte X (Cytena) to assess the viability and cell death over time. The cell death was quantified by normalizing the green object count (representing the dead cells) to cell confluency.

### ESC-GEMM and GEMM models and in vivo experiments

ES cell targeting and generation of chimeras were performed as previously described (14). BPP ES cells were targeted with TRE-shPhgdh or TRE-shRen (15) and BP ES cells were targeted with LSL-shPten-shPhgdh or LSL-shPten-shRen by recombination- mediated cassette exchanged. The latter targeting vectors were cloned by linearizing cTGME shPten (Addgene plasmid #135667) (14) using EcoRI/MluI digestion and ligating the PCR-amplified and EcoRI/MluI-digested shPhgdh or shRen fragments from TRE- shPhgdh or TRE-shRen, respectively, using primers described previously (16). The GFP- shPten-shPhgdh and GFP-shPten-shRen fragments were then PCR amplified and InFusion cloned into EcoRI-linearized cEF1a-LSL-GFP (Addgene #135672) which was modified to replace the EF1α promoter with a CMV promoter by InFusion cloning. All mouse experiments were performed under an IACUC protocol approved by the University of South Florida. Melanoma initiation was induced by applying 2.5mg/ml (BPP mice harboring TRE-shPhgdh or TRE-shRen) or 25mg/ml (BP mice harboring LSL-shPten- shPhgdh or LSL-shPten-shRen and BPP mice treated with PLX4720 chow ± Ser/Gly) of 4-Hydroxytamoxifen (Millipore Sigma, Cat. # H6278) on the shaved backs of 3-5 week old mice. For the TRE-shPhgdh/TRE-shRen experiment, shRNA expression was activated by feeding mice Doxycycline chow (200mg/kg) (Envigo, Cat. # TD180625) ad libitum. Serine/Glycine replete and Serine/Glycine free diets were purchased from Envigo (TD.110839 and TD.160752, respectively), PLX4720, and analog of PLX4032 with better bioavailability when used in chow, was purchased from Selleckchhem (Cat. # S1152) and used at 200mg/kg to formulate PLX4720-containing diets. Mice were euthanized once the tumors reached IACUC-approved endpoints or started to ulcerate. Tumors were collected for further analysis. To investigate the effect of acute BRAF^V600E^ inhibition on allograft tumors, NOD Scid Gamma (NSG) mice bearing M10M3 tumors were treated orally once daily with either vehicle or PLX4032 50mg/kg for 3 days. The tumors were collected 6 hours after the last treatment.

### Statistical Analysis

Statistical analyses were performed using GraphPad Prism 10 software. All experiments were performed at least in triplicates and each experiment was repeated at least once. A p-value of <0.05 was considered statistically significant. Survival curves were analyzed using the Log Rank (Mantel-Cox) test, while other experiments were analyzed using either Two-way ANOVA/One-way ANOVA or paired t-tests.

## RESULTS

### PHGDH expression is high in melanoma irrespective of its amplification status

To investigate the correlation of *PHGDH* expression and copy number in melanoma, we first analyzed the TCGA Skin Cutaneous Melanoma dataset. This analysis revealed that *PHGDH* amplified melanomas exhibit moderately increased *PHGDH* expression while melanomas having *PHGDH* copy number gains displayed *PHGDH* expression that was comparable to melanomas having a diploid *PHGDH* locus (**Fig. 1A**). We then analyzed the expression of *PHGDH* in melanoma using a publicly available RNA sequencing dataset comparing melanoma to nevi (GSE3189) (17). Interestingly, *PHGDH* expression is elevated in most melanomas (**Fig. 1B**). Prior studies estimate *PHGDH* amplification to occur in 10-30% of melanoma cases (3, 18) and 6% of melanomas in the TCGA-SKCM dataset exhibit *PHGDH* amplification. To analyze this further, we queried a DepMap data set on human melanoma cell lines and found a poor correlation between *PHGDH* expression and copy number status (**Fig. 1C**). We then assessed PHGDH copy number status and expression in a panel of human melanoma and melanocyte cell lines. Only two melanoma cell lines have 4 or more copies of *PHGDH* (**Fig. 1D**). However, five out of ten and nine out of ten melanoma cell lines showed elevated expression of PHGDH mRNA and protein, respectively, compared to the average expression of four melanocyte cell lines (**Fig. 1E, F, Supplementary Fig. S1A**). Accordingly, *PHGDH* copy number was only weakly correlated with PHGDH mRNA and protein expression, while PHGDH protein levels strongly correlated with *PHGDH* mRNA expression (**Fig. 1G-I**). These results indicate that PHGDH expression is frequently increased in melanoma in the absence of copy number alterations.

**Fig. 1:**
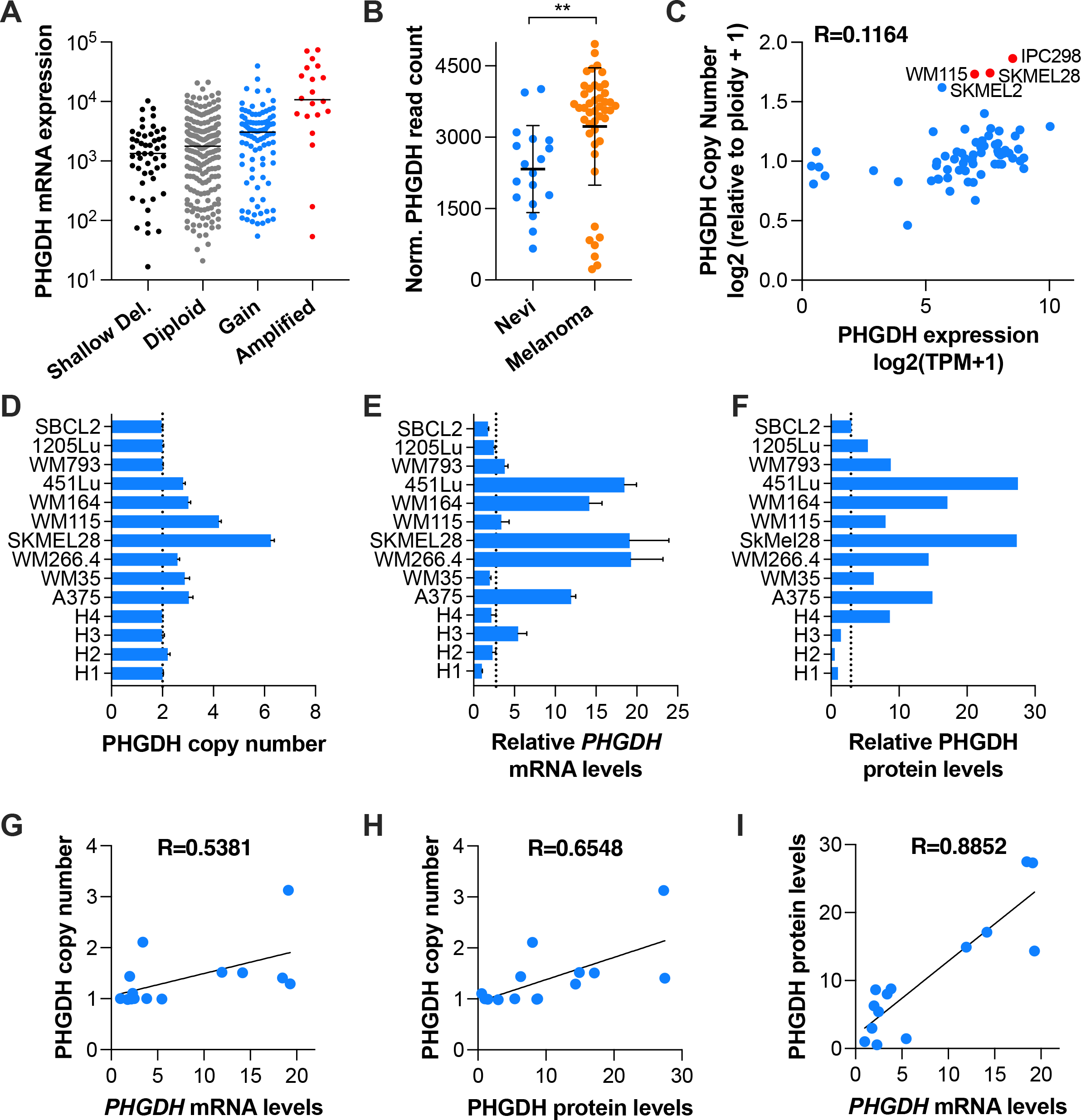
PHGDH is upregulated in melanoma irrespective of its amplification status. **(A)** *PHGDH* mRNA expression and copy number status in human melanoma tumors from the Skin Cutaneous Melanoma TCGA Pan Cancer Atlas dataset. **(B)** *PHGDH* expression comparing Nevi (n=18) vs Melanoma (n=45) in the GSE3189 dataset. **(C)** Correlation analysis between *PHGDH* mRNA expression and copy number status in melanoma cell lines (n=69) from the DepMap dataset (22Q2). The cell lines marked in red are considered amplified. **(D)** *PHGDH* absolute copy number in melanocyte and melanoma cell lines was quantified by Droplet Digital PCR using RNaseP as endogenous control. Values shown are the Mean±SD of a representative experiment performed in triplicate. H1, Hermes1; H2, Hermes2; H3, Hermes3A; H4, Hermes4B. **(E)** *PHGDH* mRNA expression in melanocyte and melanoma cell lines determined using qRT-PCR. The values are relative to Hermes1 (H1) cells and normalized using 18S as internal control. **(F)** PHGDH protein expression quantified using ImageJ software using the image from Supplementary Fig. S1A. The values shown are relative to Hermes1 (H1) cells and normalized using β-Actin. **(G-I)** Simple linear correlation analysis between *PHGDH* copy number and mRNA levels **(G)**, *PHGDH* copy number and PHGDH protein levels **(H),** and PHGDH mRNA and protein levels **(I)**. R represents the correlation coefficient. N=2 biological replicates. ** p<0.01.

### PHGDH expression and extracellular serine and glycine are important for melanoma cell proliferation in vitro

Given the increased expression, we next investigated whether PHGDH is critical for sustaining proliferation of melanoma cells in vitro. To this end, we silenced PHGDH in five non-amplified (1205Lu, WM793, WM35, WM164, A375) and one amplified (SKMel28) melanoma cell lines using two different lentiviral shRNAs (**Supplementary Fig. S1B**).

The knockdown of PHGDH significantly reduced melanoma cell proliferation in all six cell lines (**Supplementary Fig. S1C**). The increased expression of PHGDH and its important role in melanoma cell proliferation may suggest that melanoma cells rely less on uptake of extracellular serine and glycine (S/G). To investigate this, we performed proliferation assays in S/G-depleted media. Interestingly, the depletion of extracellular S/G also significantly reduced melanoma cell proliferation in all six cell lines (**Supplementary Fig. S1D**). Thus, PHGDH expression and uptake of extracellular S/G both support melanoma cell proliferation in vitro.

### PHGDH is required for melanoma development in vivo

We next examined the contribution of PHGDH expression and extracellular S/G to melanoma formation. To this end, we used our embryonic stem cell-genetically engineered mouse model (ESC-GEMM) platform (14) to generate high-contribution chimeric mice harboring oncogenic Braf (LSL-Braf^V600E^) and conditional Pten knock out (Pten^FL/FL^) alleles (BPP). High-contribution chimeras (**Supplementary Fig. S2A**) were treated with 4-Hydroxytamoxifen (4OHT) to induce melanomagenesis and fed either a control diet (+S/G diet) or an S/G-deficient diet (-S/G diet) that has previously been shown to significantly reduce systemic serine and glycine (19). Unexpectedly given the in vitro proliferation results, the S/G-deficient diet had no effect on the survival of BPP mice (**Fig. 2A**) or on the number of tumors that they developed (**Fig. 2B**). Phgdh expression was not consistently upregulated in these tumors in response to dietary S/G depletion (**Fig. 2C, Supplementary Fig. S2B**). Thus, the formation of primary melanomas is insensitive to a reduction of extracellular S/G levels, suggesting that the serine synthesis pathway may be the main source of serine. To investigate the role of PHGDH in melanoma development, we targeted BPP ES cells with a Doxycycline-inducible, GFP-linked shRNA targeting either murine Phgdh (15) or Renilla Luciferase control (20) (**Supplementary** Fig. 2C) and generated high contribution chimeras (**Supplementary Fig. S2A**). Chimeras were treated with 4OHT to activate the melanocyte-specific Cre allele and induce melanomagenesis by recombining the LSL-Braf^V600E^ and Pten conditional knockout alleles. Cre also induces the expression of the reverse Tet transactivator (rtTA3) by excising the LSL cassette from the CAG-LSL-rtTA3 allele (21). Immediately following 4OHT administration, chimeras were placed on a Dox-containing diet to induce shRNA expression in melanocytes. Notably, the silencing of Phgdh markedly prolonged the overall survival of BPP^TRE-shPten^ chimeras (**Fig. 2D**) and reduced tumor burden (**Fig. 2E**). To validate Phgdh silencing, we collected tumors at endpoint and performed Western blots. Notably, none of the shPhgdh melanomas exhibited silencing of Phgdh (**Fig. 2F, Supplementary Fig. S2D**). Based on GFP levels as a surrogate marker for shRNA expression and in contrast to shRen control shRNA, shPhgdh was not expressed (**Fig. 2F, Supplementary Fig. S2D**). Our previous studies demonstrated that global Phgdh silencing had no overt effects on the health of mice (15), indicating the Phgdh silencing is not simply inducing cell death of melanocytes. We previously observed strong selective pressure against inducible alleles that are detrimental to melanoma cells, either by transcriptional downregulation of the allele or by failed recombination of the CAG-LSL- rtTA3 allele, leading to the emergence of escaper tumors (22, 23). To circumvent this, we used an alternative approach using tandem shRNAs where one shRNA targets the gene of interest while the other is required for tumorigenesis, thereby enforcing expression of the former (24). We created constructs harboring GFP-linked shPhgdh or shRen in tandem with shPten, where Cre-mediated excision of a lox-Stop-lox cassette results in constitutive expression of the tandem shRNAs (termed LSL-shPten-shPhgdh and LSL- shPten-shRen mice) (**Supplementary Fig. S2C**). We targeted BP (LSL-Braf^V600E^; Pten^FL/WT^) ES cells with the tandem shRNA constructs, generated high-contribution chimeras (**Supplementary Fig. S2A**), and induced melanomagenesis by topical 4OHT administration. BPP^LSL-shPten-shPhgdh^ mice exhibited reduced melanoma penetrance with only two mice developing one tumor each, resulting in significantly prolonged overall survival (**Fig. 2G,H**). One of these two tumors expressed GFP – the surrogate for shRNA expression – and had reduced Phgdh levels, while the other was an escaper tumor (**Fig. 2I**). Notably, despite the reduced tumor penetrance in BPP^LSL-shPten-shPhgdh^ mice, we observed extensive formation of nevi that was comparable to BPP^LSL-shPten-shRen^ control mice (**Supplementary Fig. S2E**), indicating that Phgdh plays critical roles at the transition from nevus to frank melanoma. Thus, two different genetically engineered mouse models demonstrate that PHGDH is critical for melanoma development.

**Fig. 2:**
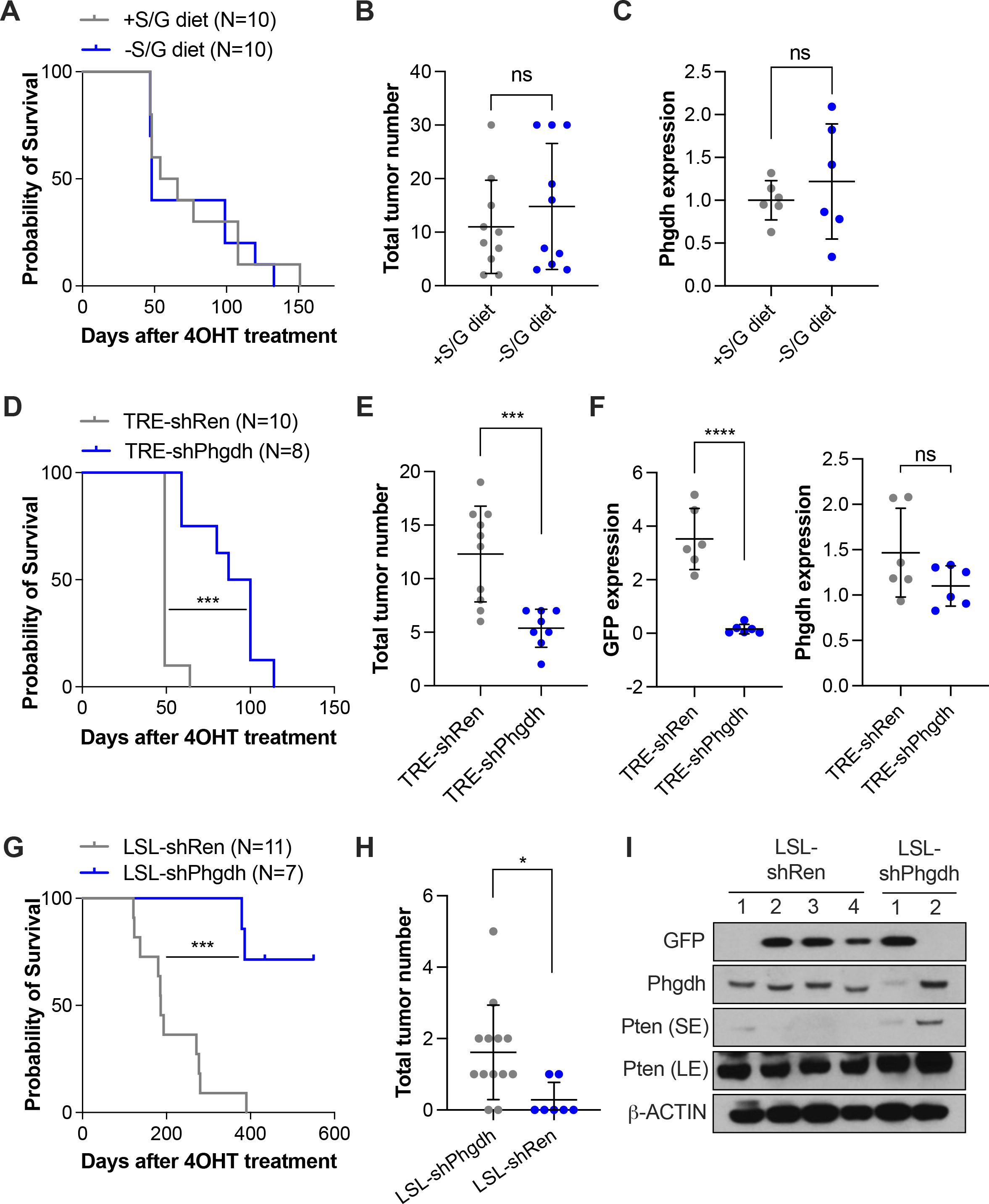
PHGDH but not extracellular S/G is critical for melanoma formation. **(A)** Kaplan Meier curve showing the overall survival of Braf^V600E^; Pten^FL/FL^ chimeras fed +S/G or -S/G diets. **(B)** Total number of tumors developed by mice shown in (A). **(C)** Phgdh expression in tumors of mice shown in (A). The quantification of Supplementary Fig. S1B is shown. **(D)** Kaplan Meier curve showing the overall survival of BPP^TRE-shPhgdh^ (TRE-shPhgdh) and BPP^TRE-shRen^ (TRE-shRen) chimeras. **(E)** Total number of tumors developed by mice shown in (D). **(F)** GFP and Phgdh expression in tumors of mice shown in (D). The quantification of Supplementary Fig. S2C is shown. **(G)** Kaplan Meier curve showing the overall survival of BP^LSL-shPten-shPhgdh^ (LSL-shPhgdh) and BP^LSL-shPten-shRen^ (LSL-shRen) chimeras. **(H)** Total number of tumors developed by mice shown in (G). **(I)** Western blot showing GFP and Phgdh expression in tumors of mice shown in (G). SE, short exposure; LE, long exposure. The survival curves were analyzed using Log-rank (Mantel-Cox) test. ****p<0.0001, ***p<0.001, *p<0.05; ns, not significant.

### PHGDH is regulated by oncogenic MAPK signaling in melanoma

Given its frequently elevated expression in melanoma, we investigated the regulation of PHGDH. Oncogenic Kras^G12D^, an upstream activator of the MAPK pathway, increased Phgdh expression in pancreatic ductal adenocarcinoma (19), and MAPK signaling is almost universally hyperactivated in melanoma. We therefore hypothesized that PHGDH is regulated by MAPK pathway signaling. We first tested whether hyperactivation of the MAPK pathway in immortalized human melanocytes (Hermes1) affects PHGDH expression. We previously established a Hermes1 cell line that constitutively expresses BRAF^V600E^ (25) and PHGDH mRNA and protein levels are increased compared to the parental Hermes1 cells (**Supplementary** Fig. 3A,B). Similarly, using a Dox-inducible vector, acute expression of BRAF^V600E^ for 72 hours increased PHGDH mRNA and protein levels in Hermes1 parental cells (**Fig. 3A,B**). We then analyzed PHGDH expression upon BRAF^V600E^ inhibition with Vemurafenib (PLX4032). Inhibition of oncogenic BRAF in 1205Lu human melanoma cells suppressed PHGDH mRNA and protein expression in a time-dependent manner, with maximal downregulation after 72 hours (**Fig. 3C,D, Supplementary Fig. S3C,D**). Inhibition of MEK1/2 with Selumetinib (AZD6244) or ERK1/2 with SCH772984 also reduced PHGDH mRNA and protein expression in 1205Lu cells after 72 hours (**Fig. 3C,D**). Similar effects of MAPK pathway inhibition on Phgdh expression were observed in the murine melanoma cell line M10M3 (**Fig. 3E, F**). Additionally, inhibition of the MAPK pathway downregulated PHGDH expression in the *PHGDH*-amplified human melanoma cells SKMel28 (**Supplementary Fig. S3E,F**). To investigate if MAPK pathway inhibition reduces PHGDH expression in vivo, we established allograft tumors using murine M10M3 cells in NSG mice. We treated tumor-bearing mice orally with 50mg/kg PLX4032 daily for 3 days, collected the tumors, and determined Phgdh expression. In line with our in vitro observations, PLX4032 administration reduced PHGDH protein levels in allograft tumors (**Fig. 3G, Supplementary Fig. S3G**). These results demonstrate that the MAPK pathway regulates PHGDH expression in melanoma.

**Fig. 3:**
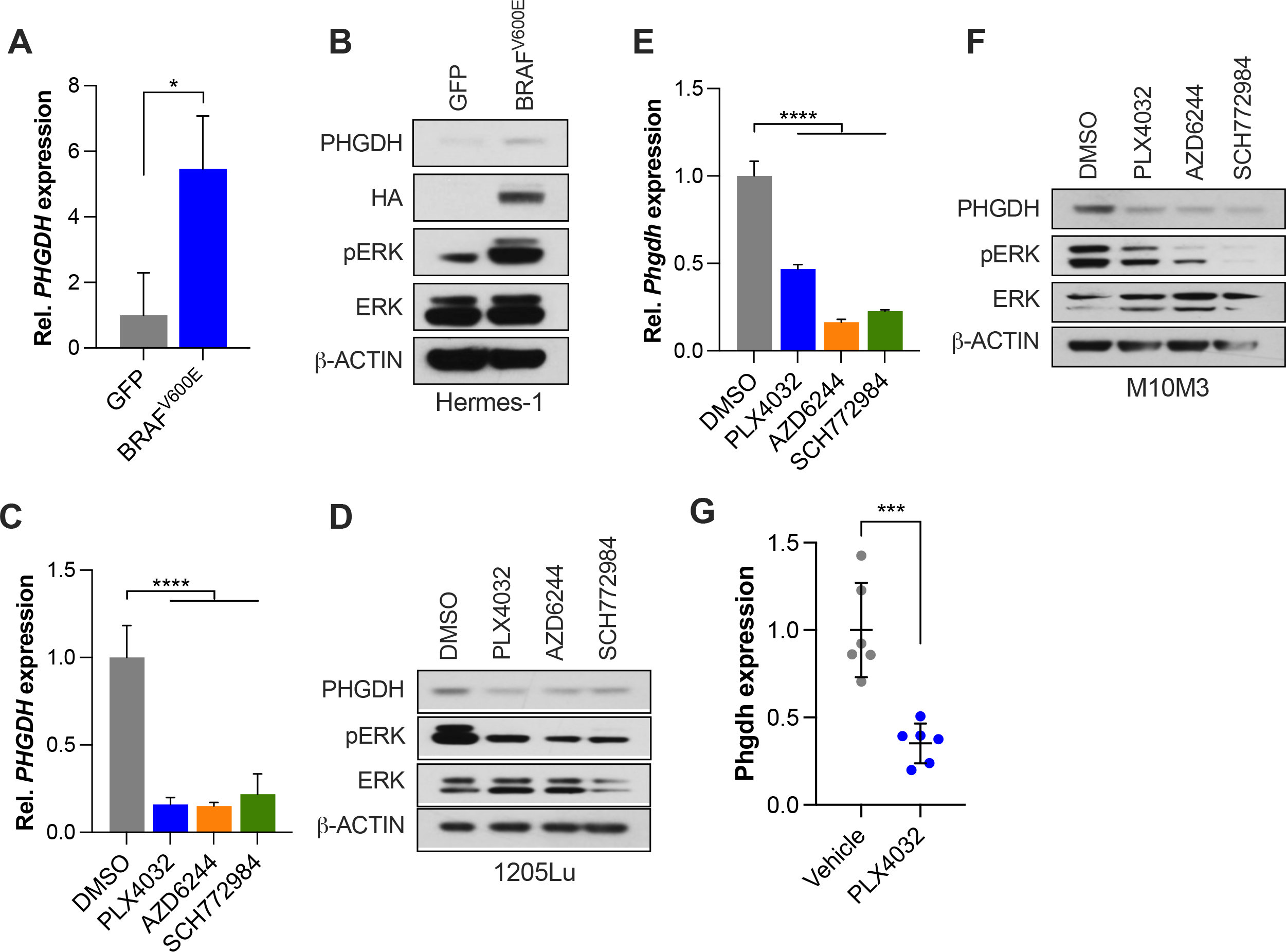
Mutant BRAF^V600E^ promotes PHGDH expression. **(A, B)** PHGDH mRNA (A) and protein (B) expression upon acute overexpression of mutant BRAF^V600E^ in Hermes1 melanocytes. BRAF^V600E^ expression was induced by 0.5μg/ml Doxycycline for 72 hours. **(C, D)** PHGDH mRNA (C) and protein (D) expression in response to MAPK pathway inhibition using BRAF^V600E^ inhibitor (PLX4032), MEK inhibitor (AZD6244), or ERK inhibitor (SCH772984) at 0.25μM in human 1205Lu cells after 72 hours. **(E, F)** Phgdh mRNA (E) and protein (F) expression in response to MAPK pathway inhibition using BRAF^V600E^ inhibitor (PLX4032), MEK inhibitor (AZD6244), or ERK inhibitor (SCH772984) at 0.25μM in murine M10M3 cells after 72 hours. **(G)** Quantification of PHGDH expression in M10M3 allograft tumors after acute treatment with PLX4032 orally once daily for 3 days. Tumors were collected 6 hours after last dose. The quantification of Supplementary Fig. S3G is shown. ****p<0.0001, *p<0.05; ns, not significant.

### The MAPK pathway regulates PHGDH via an mTOR-ATF4 axis

The MAPK pathway induces the transcription factor ATF4 (26–28), which is a transcriptional activator of *PHGDH* expression (29, 30). Moreover, ATF4 translation is regulated by mTORC1 (31, 32) and oncogenic BRAF promotes mTORC1 activity via ERK-mediated inhibitory phosphorylation of TSC2 (33–36). We therefore tested whether MAPK signaling regulates PHGDH expression via an mTORC1/ATF4 axis. Inhibition of the MAPK pathway for 72 hours significantly suppressed ATF4 protein levels in 1205Lu and SKMel28 cells with concomitant decreases of pTSC2 and the mTORC1 effectors pS6 and p4EBP1 (**Fig. 4A**, **Supplementary Fig. S4A**). MAPK inhibition also reduced ATF4 expression in M10M3 cells even though the inhibitors only decreased p4EBP1 but not pS6 (**Fig. 4B**). Expression of mutant BRAF^V600E^ increased the levels of pTSC2, pS6, p4EBP1, and ATF4 in melanocytes (**Fig. 4C**). Pharmacological inhibition of mTORC1 with Temsirolimus for 72 hours reduced the mRNA and protein levels of PHGDH and this was associated with reduced ATF4 expression (**Fig. 4D,E, Supplementary Fig. S4B,C**). Additionally, silencing of ATF4 using pooled siRNAs decreased PHGDH mRNA and protein expression (**Fig. 4F,G**, **Supplementary Fig. S4D,E**). These results suggest that the MAPK pathway controls PHGDH expression via mTORC1-mediated regulation of ATF4.

**Fig. 4:**
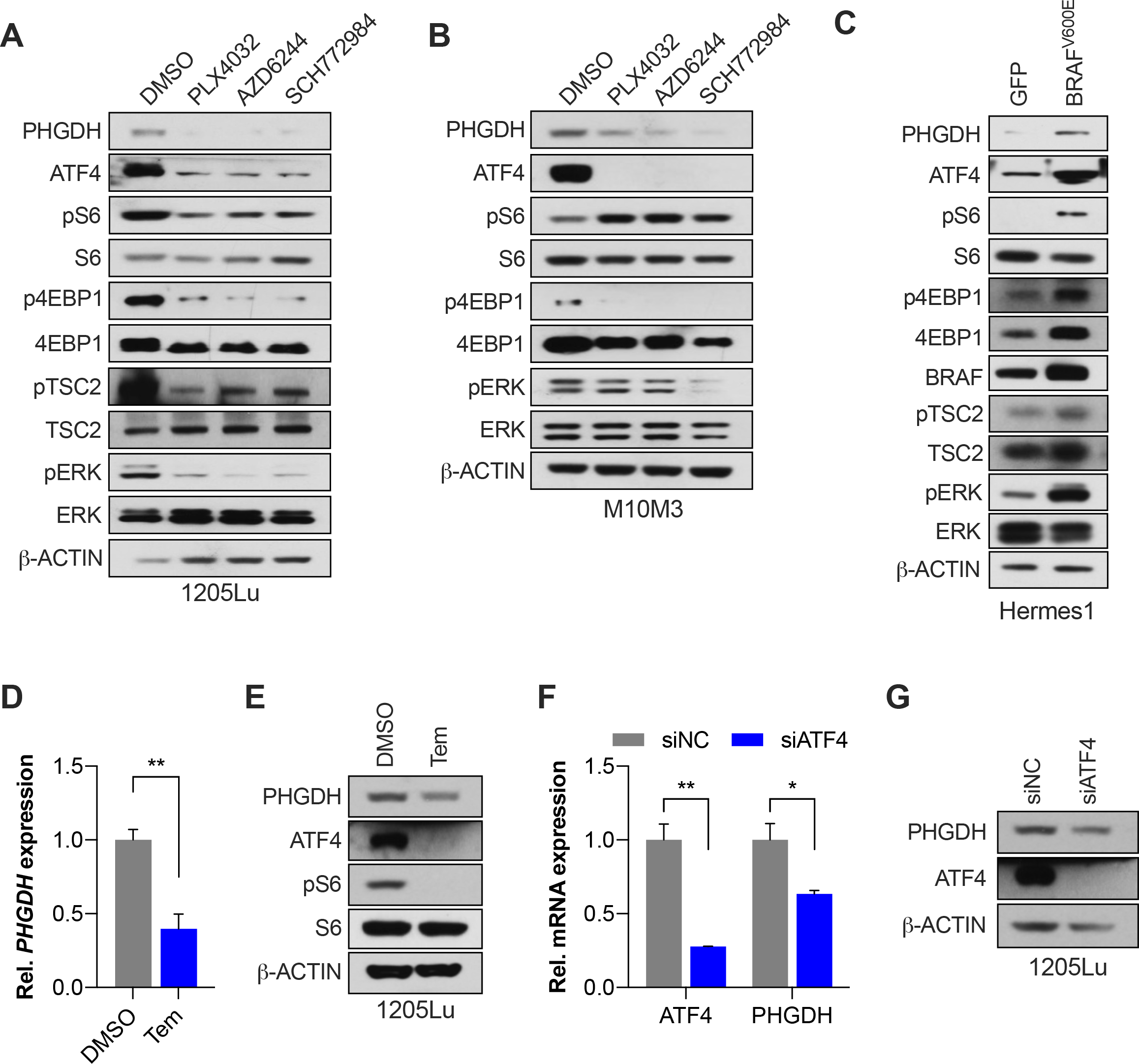
BRAFV600E promotes PHGDH expression via the mTORC1-ATF4 axis. **(A, B)** Western blot showing the effect of treatment with 0.25μM PLX4032, AZD6244, or SCH772984 for 72 hours on PHGDH, ATF4, pTSC2, pS6, and p4EBP1 expression in 1205Lu cells (A) and M10M3 cells (B). **(C)** Western blot showing the effect of acute overexpression of BRAF^V600E^ on PHGDH, ATF4, pTSC2, pS6, and p4EBP1 expression in Hermes1 cells after 72 hours of 0.5μg/ml Doxycycline. **(D, E)** *PHGDH* mRNA (D) and PHGDH and ATF4 protein expression (E) in 1205Lu cells in response to 0.1μM Temsirolimus treatment for 72 hours. **(F, G)** PHGDH and ATF4 mRNA (F) and protein (G) expression in 1205Lu cells upon *ATF4* silencing using siRNA. Tem, Temsirolimus. **p<0.005, *p<0.05.

### The MAPK pathway regulates the translation of ATF4

mTOR regulates the translation of ATF4 via 4EBPs (31) and we investigated whether the MAPK pathway influences ATF4 levels through mTORC1-mediated translation. We used an ATF4 translation reporter where the upstream open reading frames (uORFs) required for cap-dependent translation of ATF4 are linked to GFP (10). We treated 1205Lu and M10M3 cells stably expressing the ATF4 translation reporter with Vemurafenib for 48 hours and observed robust downregulation of GFP reporter expression as well as endogenous ATF4 expression (**Fig. 5A, Supplementary Fig. S5A**). Temsirolimus treatment similarly reduced reporter and endogenous ATF4 expression (**Fig. 5A, Supplementary Fig. S5A**). ATF4 translation is also induced by the integrated stress response (ISR) pathway upon amino acid starvation where uncharged tRNAs activate GCN2-eIF2α signaling (37). However, amino acid depletion, including serine starvation (38), lowers mTORC1 activity. We therefore tested whether serine starvation induces ATF4 translation in melanoma cells, whether this is mediated by mTORC1 and/or the ISR, and whether the MAPK pathway contributes under these conditions. Serine starvation robustly induced ATF4 expression in a panel of human melanoma cell lines with a concomitant moderate increase in PHGDH levels (**Supplementary Fig. S5B**). In 1205Lu cells, MAPK and mTORC1 inhibition, but not GNC2 inhibition, reduced reporter and endogenous ATF4 expression also under serine starvation (**Fig. 5A**). Conversely, inhibition of MAPK, GCN2, and to a lower extent mTORC1 reduced reporter and endogenous ATF4 expression in serine-starved M10M3 cells (**Supplementary Fig. S5A**). Thus, while the contribution of the ISR to ATF4 translation under basal and serine restricted conditions may be cell line- and/or species-dependent, MAPK signaling and mTORC1 are potent inducers of ATF4 translation.

**Fig. 5:**
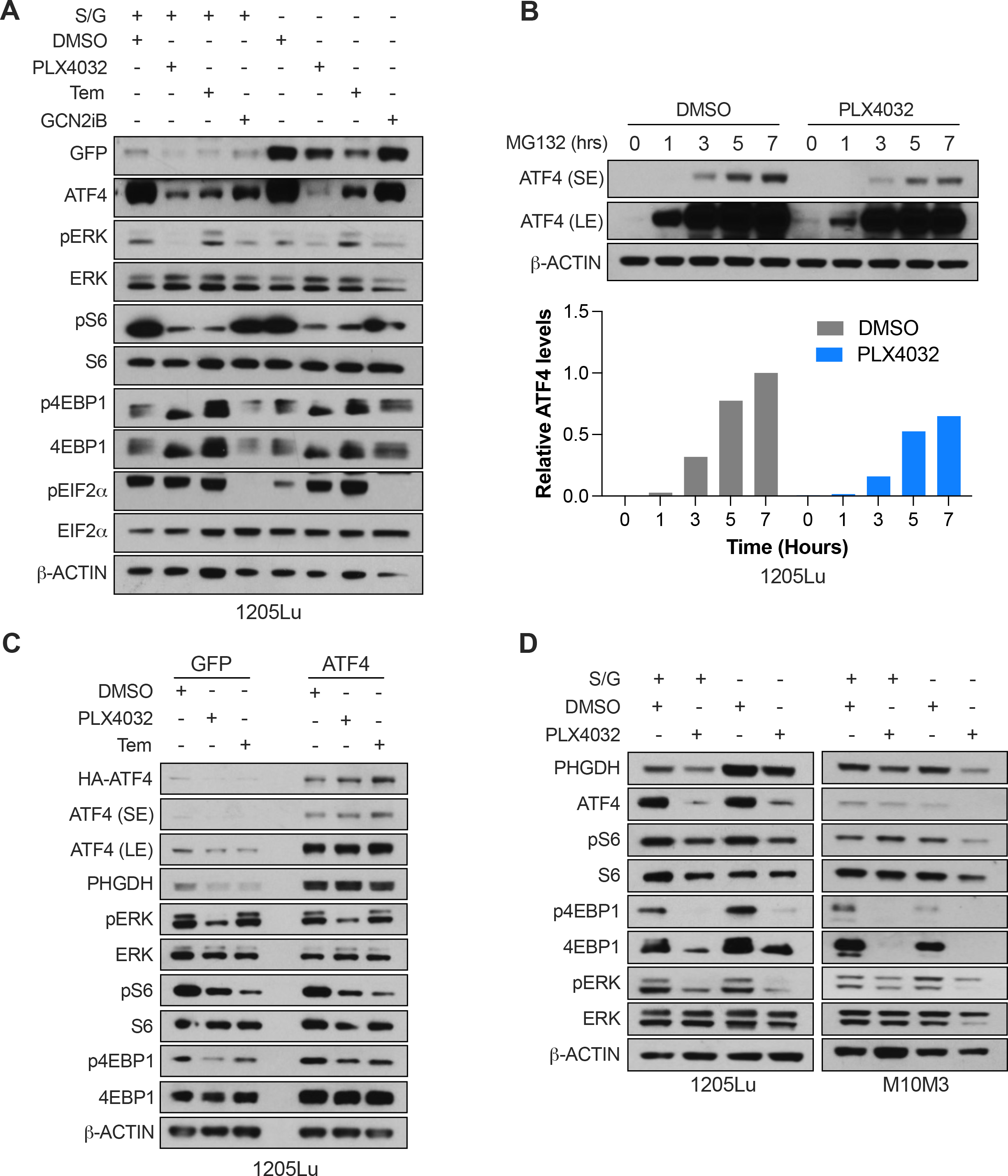
The MAPK pathway regulates the translation of ATF4. **(A)** Western blot showing the effect of 48 hours of treatment with PLX4032 (0.25μM), Temsirolimus (Tem, 0.1μM), or the GNC2 inhibitor GCN2iB (20μM) on the expression of the uORF-ATF4-GFP reporter, endogenous ATF4, and pEIF2α in 1205Lu cells grown in +S/G or -S/G medium. **(B)** Western blot and quantification showing the effect of 48 hours of PLX4032 treatment (0.25μM) on ATF4 accumulation upon proteasome inhibition with MG132 (10μg/ml) in 1205Lu cells. **(C)** Western blot showing the effect of 72 hours of PLX4032 (0.25μM) or Temsirolimus (0.1μM) treatment on the doxycycline inducible expression of endogenous ATF4 and mTORC1-independent ectopic HA-ATF4 in 1205Lu cells. **(D)** Western blot showing the effect of combined PLX4032 treatment (0.25μM) for 72 hours and S/G depletion on PHGDH and ATF4 protein levels in 1205Lu and M10M3 cells.

To further investigate the kinetics of ATF4 translation in 1205Lu and M10M3 cells, we inhibited the proteosome using MG132 in the presence or absence of Vemurafenib. ATF4 rapidly accumulated in DMSO-treated control cells upon MG132 administration, and this accumulation was significantly diminished in the presence of Vemurafenib (**Fig. 5B, Supplementary Fig. S5C**). We then expressed HA-tagged ATF4 in 1205Lu cells and treated with Vemurafenib. This ectopic construct does not contain the ATF4 uORFs rendering the translation of ectopic HA-ATF4 independent of mTORC1, which was confirmed by the lack of an effect of Temsirolimus on the expression of ectopic HA-ATF4 (**Fig. 5C**). Similarly, Vemurafenib treatment had no effect on ectopic HA-ATF4 expression and, importantly, expression of ectopic HA-ATF4 rescued the reduction of PHGDH expression in response to Vemurafenib (**Fig. 5C**). These results indicate that MAPK signaling augments the ATF4-PHGDH axis by promoting ATF4 translation.

### Inhibition of BRAF^V600E^ sensitizes melanoma cells to serine/glycine starvation

Our data showed that despite constitutive MAPK hyperactivation, melanoma cells increase PHGDH expression upon S/G starvation (**Supplementary Fig. S5B**). We tested if this response is independent of the MAPK pathway. We starved 1205Lu, SKMel28, and M10M3 cells of S/G for 72 hours with or without the treatment with Vemurafenib. Surprisingly, while S/G starvation increased PHGDH levels in the control condition, this increase was significantly diminished by Vemurafenib (**Fig. 5D, Supplementary Fig. S5D**). Moreover, unlike what has been observed in HCT116 cells (38), mTORC1 activity was maintained during S/G starvation given the unchanged levels of pS6 and p4EBP1 (**Fig. 5D, Supplementary Fig. S5D**). Thus, MAPK activity is required for melanoma cells to respond to scarce extracellular serine levels, possibly via the mTORC1-ATF4 axis.

The above finding led us to hypothesize that inhibition of BRAF^V600E^ sensitizes melanoma cells to serine starvation by attenuating the increase in PHGDH expression. We treated 1205Lu and A375 melanoma cells with Vemurafenib for 72 hours in regular or S/G-depleted media and determined uptake of the SYTOX Green dead cell dye. Vemurafenib induced cell death in both cell lines, while S/G depletion also caused some cell death in A375 cells (**Fig. 6A, Supplementary Fig. S6A**). However, the combination of Vemurafenib and S/G starvation led to a significant increase in melanoma cell death (**Fig. 6A, Supplementary Fig. S6A**). To investigate the effect of Vemurafenib combined with S/G starvation on long-term melanoma cell survival, we performed clonogenic cell survival assays with 1205Lu, A375, and SKMel28 cells. Individual treatments of Vemurafenib or S/G starvation significantly reduced the growth of all three melanoma cell lines and combining Vemurafenib with S/G starvation had an even more pronounced effect (**Fig. 6B, Supplementary Fig. S6B**). These results indicate that the inhibition of mutant BRAF^V600E^ sensitizes melanoma cells to extracellular S/G depletion.

**Fig. 6:**
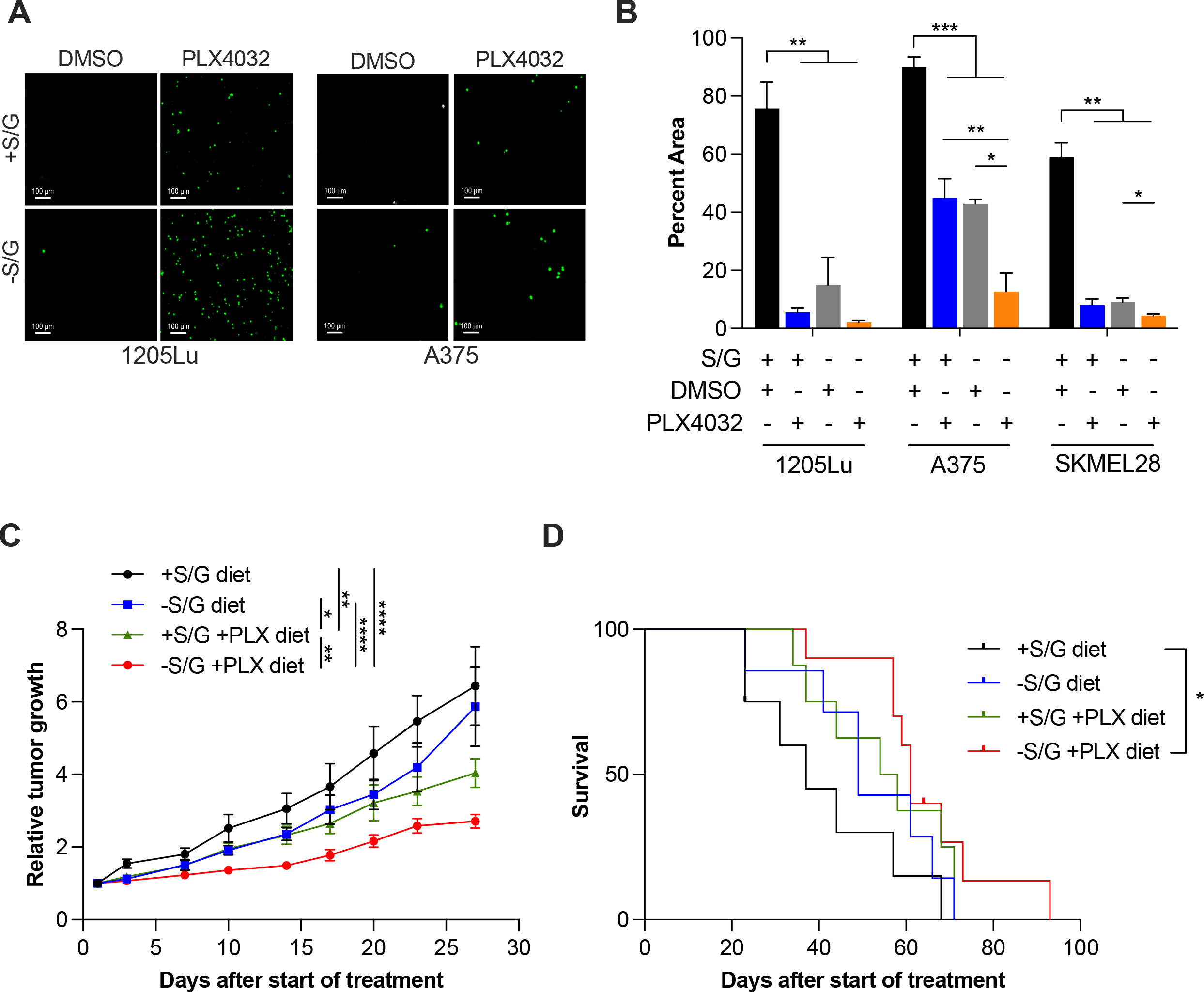
BRAF inhibition sensitizes melanoma to S/G starvation. **(A)** Images showing the effect of S/G starvation and PLX4032 treatment on cell death for 72 hours in 1205Lu (0.25μM) and A375 (0.5μM) cells. Dying and dead cells are labeled with the SYTOX green dye. **(B)** Quantification of colony growth assays of cells treated with PLX4032 (1205Lu, 0.25μM; A375, 0.5μM; SKMel28, 1μM) and grown in +S/G or - S/G medium for 10-15 days. **(C)** Graph showing the effect of different diets on melanoma growth in Braf^V600E^, Pten^FL/FL^ mice. Tumor growth was normalized to the tumor size at the start of treatment. **(D)** Kaplan Meier curve showing the overall survival of Braf^V600E^, Pten^FL/FL^ mice treated on diets with or without S/G and with or without PLX4720.

### Vemurafenib sensitizes melanoma to dietary S/G restriction

We next investigated the effect of combining BRAF^V600E^ inhibition with dietary S/G restriction in vivo. To this end, we induced melanomagenesis in conventionally bred LSL- Braf^V600E^; Pten^FL/FL^; Tyr-CreERt2 (BPP) mice with 4OHT. Once palpable tumors (∼100 mm3) developed, BPP mice were randomly switched to one of four diets: a S/G-containing control diet, a S/G-depleted diet, a diet containing a moderate dose of Vemurafenib (200mg/kg), or a S/G-depleted diet containing 200mg/kg Vemurafenib. Monitoring melanoma volumes over time revealed that the reduction in tumor size elicited by Vemurafenib treatment was significantly enhanced by the combination of Vemurafenib and dietary S/G restriction (**Fig. 6C, Supplementary Fig. S6C**). The reduction in tumor growth in BPP mice receiving the S/G-deficient diet containing Vemurafenib resulted in a moderate but significant extension of overall survival while either treatment alone had no significant effect on survival (**Fig. 6D**). We harvested melanomas at endpoint from the four cohorts and analyzed the expression of Phgdh and pERK by immunohistochemistry. In line with our in vitro observations, Phgdh expression was decreased in tumors from mice treated with Vemurafenib both in the presence or absence of dietary S/G and this correlated with reduced pERK staining (**Supplementary Fig. S6D**). These results demonstrate that dietary S/G restriction enhances the effect of Vemurafenib on melanoma in vivo.

## DISCUSSION

PHGDH overexpression is induced in and promotes melanoma progression (5, 18), metastasis (7), and therapy resistance (8, 9). However, it has remained unknown whether PHGDH is required for melanoma formation. This study demonstrates that PHGDH expression is universally upregulated in melanoma via BRAF^V600E^-induced ATF4 translation, rendering melanoma formation critically dependent upon PHGDH. This creates a vulnerability whereby PHGDH expression can be diminished in BRAF^V600E^ mutant melanoma using targeted therapies such as Vemurafenib, thereby sensitizing melanoma cells to extracellular serine restriction.

This study provides the first genetic evidence for a critical role of PHGDH in the formation of melanoma. Together with the genetic depletion and pharmacologic inhibition of PHGDH in progression, metastasis, and resistance (5, 7–9, 18), our results indicate that PHGDH is important throughout melanomagenesis. In addition to melanoma, PHGDH undergoes recurrent copy number gains in breast cancer (3, 4) and is overexpressed in multiple cancer types (3, 4, 18, 39–47). RNAi-mediated depletion and pharmacologic inhibition indicate that PHGDH supports xenograft tumor growth of various cancer types including lung (10, 48), breast (49), pancreatic (50), thyroid (51), colon (52, 53), esophageal (54), endometrial (55), prostate (56), neuroblastoma (57), and Ewing sarcoma (41, 58). However, whether PHGDH is essential for the initiation and/or formation of these cancers has not been addressed in genetic mouse models. Our approach in melanoma models provides a framework with which the role of PHGDH in other cancer types may be elucidated. This will be particularly interesting for cancer types that are driven by MAPK pathway hyperactivation, such as those harboring activating mutations in KRAS. Prior studies have shown that Kras^G12D^-mutant mouse models of pancreatic adenocarcinoma (PDAC) are insensitive to dietary serine restriction while models of APC loss-of-function-driven colon cancer and Eµ-myc-driven lymphoma display extended survival on a serine-deficient diet (19). Moreover, expression of Kras^G12D^enhances the expression of the serine synthesis pathway enzymes Phgdh, Psat1, and Psph in murine PDAC cell lines (19). These data suggest that PHGDH expression is increased in other cancers exhibiting MAPK pathway hyperactivation similar to BRAF^V600E^ mutant melanoma. However, whether transcriptional control via mTORC1/ATF4 or other mechanisms regulate PHGDH expression in such cancers remains to be determined.

We demonstrate that BRAF^V600E^ increases ATF4 expression via mTORC1- mediated translational control, which in turn leads to *PHGDH* transcription. BRAF^V600E^ and the MAPK pathway have previously been shown to activate mTORC1 through RSK-mediated inhibitory phosphorylation of TSC2 (33, 36), and we confirm that MAPK inhibition reduces the levels of pTSC2 as well as the mTORC1 effectors pS6 and p4E- BP1. ATF4 is generally considered a stress-responsive transcription factor that is activated by the integrated stress response (ISR) pathway via GNC2-mediated eIF2α phosphorylation and selective translation of the ATF4 mRNA. However, ATF4 has recently been uncovered as a mTORC1 effector that mediates pro-growth signals (29, 32, 59). mTORC1 also promotes ATF4 translation, but this is independent of the ISR and eIF2α phosphorylation (31, 59). Our results demonstrate that BRAF^V600E^ controls ATF4 levels via mTORC1-mediated translation, with limited contribution by the ISR. Interestingly, serine starvation of melanoma cells only modestly increased PHGDH levels, suggesting that the BRAF^V600E^-mTORC1 axis already potently stimulates PHGDH expression in unstressed cells. However, serine starvation robustly promoted ATF4 levels in the absence of potent ISR pathway induction. Thus, ATF4 may be responsive to serine starvation in melanoma cells independent of the ISR. This response may be through mTORC1, since mTORC1 is not inactivated upon serine starvation in melanoma cells. This contrasts with other cancer types that lack BRAF mutations (35, 53), suggesting that MAPK hyperactivation maintains mTORC1 activity even under amino acid starvation. Indeed, BRAF inhibition blocks the serine starvation-mediated induction of ATF4. However, whether mTORC1 can sense and respond to amino acid starvation in BRAF^V600E^-mutant cells remains to be determined.

The MAPK pathway may regulate the levels of ATF4 through additional mechanisms. During the development of BRAF inhibitor resistance of melanoma cells, ERK phosphorylates ATF4, resulting in increased protein stability (60). This mechanism may not at play in treatment naïve melanoma cells since the ectopic expression of ATF4, which is independent of uORF-mediated translational control (i.e., translation independent of mTORC1) was unaffected upon BRAF and mTORC1 inhibition. Moreover, in lung adenocarcinoma, the MAPK pathway enhances the expression of NRF2 (61), which in turn increases ATF4 transcription (10). Additionally, the MAPK pathway promotes MYC activity in many cancers including melanoma (62, 63), which promotes ATF4 translation via GCN2 (64). Our findings add a novel mechanism of ATF4 regulation in BRAF^V600E^-mutant melanoma cells where ATF4 translation is induced via MAPK-mediated stimulation of mTORC1 activity. Thus, the regulation of ATF4 appears context and cell-type dependent.

Using a genetic approach, we showed that PHGDH plays a critical role in the progression of nevi to frank melanoma. Whether PHGDH is similarly important in established melanomas remains to be determined with more sophisticated genetically engineered mouse models. However, the upregulation of PHGDH in melanoma progression, brain metastasis survival (7), and resistance to MAPK pathway inhibition (8, 9) suggests important PHGDH functions at different stages of melanomagenesis. PHGDH has been considered as a therapeutic target for cancer therapy, and several inhibitors of PHGDH have been developed (49, 65–68). While first-generation PHGDH inhibitors suffered from poor potencies in vivo, second generation inhibitors exhibit improved efficacy profiles (53, 68). However, no clinical data are available yet that demonstrate the therapeutic utility of PHGDH inhibitors in cancer patients. Studies using colon cancer xenografts showed that a second generation PHGDH inhibitor, PH755, had minor effects on tumor growth (53), suggesting limited efficacy of the inhibitor, non-enzymatic functions of the PHGDH protein, or compensation through uptake of extracellular serine. Indeed, the combination of PHGDH inhibition with dietary serine and glycine restriction robustly impaired xenograft tumor growth (53). Interestingly, PHGDH inhibition also impaired metastasis of triple-negative breast cancer and renal cell carcinoma to the brain where extracellular serine is limited (7), indicating that combining PHGDH inhibition with extracellular serine restriction is a viable therapeutic approach. However, the combination of PH755 with dietary serine/glycine restriction resulted in significant weight loss of non-tumor bearing mice (53), suggesting that this combination may be hampered by toxicities due to the systemic lowering of serine in all tissues.

PHGDH levels in BRAF^V600E^ mutant melanoma cells modestly increase in response to serine restriction. Importantly, however, PHGDH induction by serine starvation was robustly diminished by Vemurafenib. This offered an alternative to a treatment strategy involving PHGDH inhibitors, where PHGDH levels can be maintained at low levels by BRAF^V600E^ inhibition, thus sensitizing melanoma cells to serine starvation. Encouragingly, Vemurafenib combined with serine/glycine restriction resulted in increased melanoma cell death in vitro, reduced tumor growth and extended survival in a spontaneous melanoma mouse model. The advantage of this treatment strategy is that Vemurafenib inhibits the V600E-mutant form of BRAF with high selectivity, thus reducing PHGDH levels only in melanoma cells while sparing BRAF wildtype cells. Moreover, BRAF^V600E^ inhibitors are already FDA approved and widely used to treat melanoma patients, so this combination could be readily tested in patients. Several strategies could be envisioned to starve melanomas of serine. Patients could be provided a specially formulated diet lacking serine and glycine; however, adherence to the diet may be a concern requiring pharmacological approaches. This could be achieved using enzyme-based therapy to systemically deplete extracellular serine. This approach has been very successful in T-ALL for targeting of asparagine (69), and preclinical studies targeting cysteine and arginine for cancer therapy are promising (70–72). Additionally, serine uptake could be prevented by inhibiting serine transporters such as ASCT2 (73). Thus, our findings that PHGDH is essential for melanomagenesis and regulated by BRAF^V600E^ revealed a potential treatment strategy for patients with BRAF^V600E^-mutant melanoma.

## ACKNOWLEDGMENTS

We are grateful to Karreth lab members for helpful discussions. This work was supported in part by a Miles for Moffitt Award (FAK), a Harry J. Lloyd Charitable Trust Career Development Award (FAK), a Bankhead-Coley Grant from the Florida Department of Health (FAK), and a NIH/NCI grant (P01CA250984 to FAK and GMD). This work was also supported by the Gene Targeting Core, the Proteomics Core, and the Biostatistics and Bioinformatics Shared Resource, which are funded in part by Moffitt’s Cancer Center Support Grant (P30CA076292).

## AUTHOR CONTRIBUTIONS

Conceptualization – NJ, FAK; Formal Analysis – NJ, XX, YK, KYT, GMD, FAK; Funding Acquisition – GMD, FAK; Investigation – NJ, XX, BP, YK, OV, KYT, GMD, FAK; Methodology – NJ, XX, YK, GMD, FAK; Project administration – FAK; Supervision – GMD, FAK; Visualization – NJ, FAK; Writing – original draft – NJ, FAK; Writing – review & editing – NJ, XX, BP, YK, OV, KYT, GMD, FAK.

## SUPPLEMENTARY FIGURE LEGENDS

Supplementary Fig. S1: PHGDH and extracellular S/G are important for melanoma cell proliferation in vitro.

**(A)** Western blot showing the expression of PHGDH in melanocytes and melanoma cell lines. **(B)** Western blot showing the knockdown of PHGDH using two different shRNAs (#1 and #2) compared to scrambled control (shSCR) in six melanoma cell lines. **(C)** Effect of PHGDH knockdown on melanoma cell proliferation in six melanoma cell lines. **(D)** Effect of S/G depletion on melanoma cell proliferation in six melanoma cell lines. Mean±SD of triplicates normalized to day 1 are shown. The statistical significance shown in (C) compares shSCR vs shPHGDH#1 and shSCR vs shPHGDH#2. ****p<0.0001, *** p<0.0005, *p<0.05.

Supplementary Fig. S2: Quantification of chimerism and Phgdh expression in the different mouse models.

**(A)** Percentage of chimerism of mice shown in Fig. 2A,D,G. **(B)** Western blot showing Phgdh expression in tumors from Braf^V600E^; Pten^FL/FL^ chimeras on +S/G or -S/G diets. **(C)** Western blot showing GFP and Phgdh expression in tumors from BPP^TRE-shPhgdh^ (TRE- shPhgdh) and BPP^TRE-shRen^ (TRE-shRen) chimeras. **(D)** Outline of the ESC-GEMM approach and scheme depicting the alleles used for melanocyte-specific Phgdh silencing.

Supplementary Fig. S3: PHGDH is upregulated by mutant BRAF^V600E^.

**(A,B)** PHGDH mRNA (C) and protein (D) expression upon stable overexpression of mutant BRAF^V600E^ in Hermes1 melanocytes compared to parental Hermes1 cells. **(C, D)** Time course of PHGDH mRNA (C) and protein (D) expression in 1205Lu cells in response to treatment with 0.25μM BRAF^V600E^ inhibitor (PLX4032). SE, short exposure; LE, long exposure. **(E, F)** PHGDH mRNA (E) and protein (F) expression in response to MAPK pathway inhibition for 72 hours using PLX4032 (1µM), AZD6244 (0.5µM), or SCH772984 (0.5µM) in human SKMel28 cells. **(G)** Western blot showing Phgdh expression in M10M3 allograft tumors after acute treatment with PLX4032 (50mg/kg) orally once daily for 3 days. ****p<0.0001, **p<0.01, *p<0.05; ns, not significant.

Supplementary Fig. S4: PHGDH is upregulated by BRAF^V600E^ via the mTORC1-ATF4 axis.

**(A)** Western blot showing the effect on PHGDH, ATF4, pTSC2, pS6, and p4EBP1 expression in response to MAPK pathway inhibition for 72 hours using PLX4032 (1µM), AZD6244 (0.5µM), or SCH772984 (0.5µM) in human SKMel28 cells. **(B, C)** *PHGDH* mRNA (B) and PHGDH and ATF4 protein expression (C) in response to 0.1μM Temsirolimus treatment in M10M3 and SKMEL28 cells for 72 hours. **(D, E)** PHGDH and ATF4 mRNA (D) and protein expression (E) after *ATF4* silencing using siRNA in SKMel28 cells. **p<0.005, *p<0.05

Supplementary Fig. S5: The MAPK pathway regulates the translation ATF4.

**(A)** Western blot showing the effect of 48 hours of treatment with PLX4032 (0.25μM), Temsirolimus (Tem, 0.1μM), or the GNC2 inhibitor GCN2iB (20μM) on the expression of the uORF-ATF4-GFP reporter, endogenous ATF4, and pEIF2α in M10M3 cells grown in

+S/G or -S/G medium. **(B)** Western blot showing the effect of S/G starvation on ATF4 and PHGDH expression in six human melanoma cell lines **(C)** Western blot and quantification showing the effect of 48 hours of PLX4032 treatment (0.25μM) on ATF4 accumulation upon proteasome inhibition with MG132 (10μg/ml) in M10M3 cells. **(D)** Western blot showing the effect of combined PLX4032 treatment (1μM) for 72 hours and S/G depletion on PHGDH and ATF4 protein levels in in SKMel28 cells.

Supplementary Fig. S6: S/G starvation potentiates the effect of BRAF inhibition.

**(A)** Quantification of dead 1205Lu and A375 cells in cultured in +S/G or -S/G medium and treated with PLX4032 (1205Lu, 0.25μM; A375, 0.5μM) for 72hrs. **(B)** Images of crystal violet stained colony growth assays cultured in +S/G or -S/G medium and treated with PLX4032 (1205Lu, 0.25μM; A375, 0.5μM; SKMel28, 1μM). **(C)** Growth curve of individual melanoma tumors in Braf^V600E^, Pten^FL/FL^ mice on diets with or without S/G and with or without PLX4720. **(D)** Images of immunohistochemistry stainings of Phgdh and pErk in tumors from Braf^V600E^, Pten^FL/FL^ mice on diets with or without S/G and with or without PLX4720.

## Notes

### Competing Interest Statement

The authors have declared no competing interest.

