## Supplementary figures and images for "MAPK-mediated PHGDH induction is essential for melanoma formation and represents an actionable vulnerability"

### Supplemental Figure 1

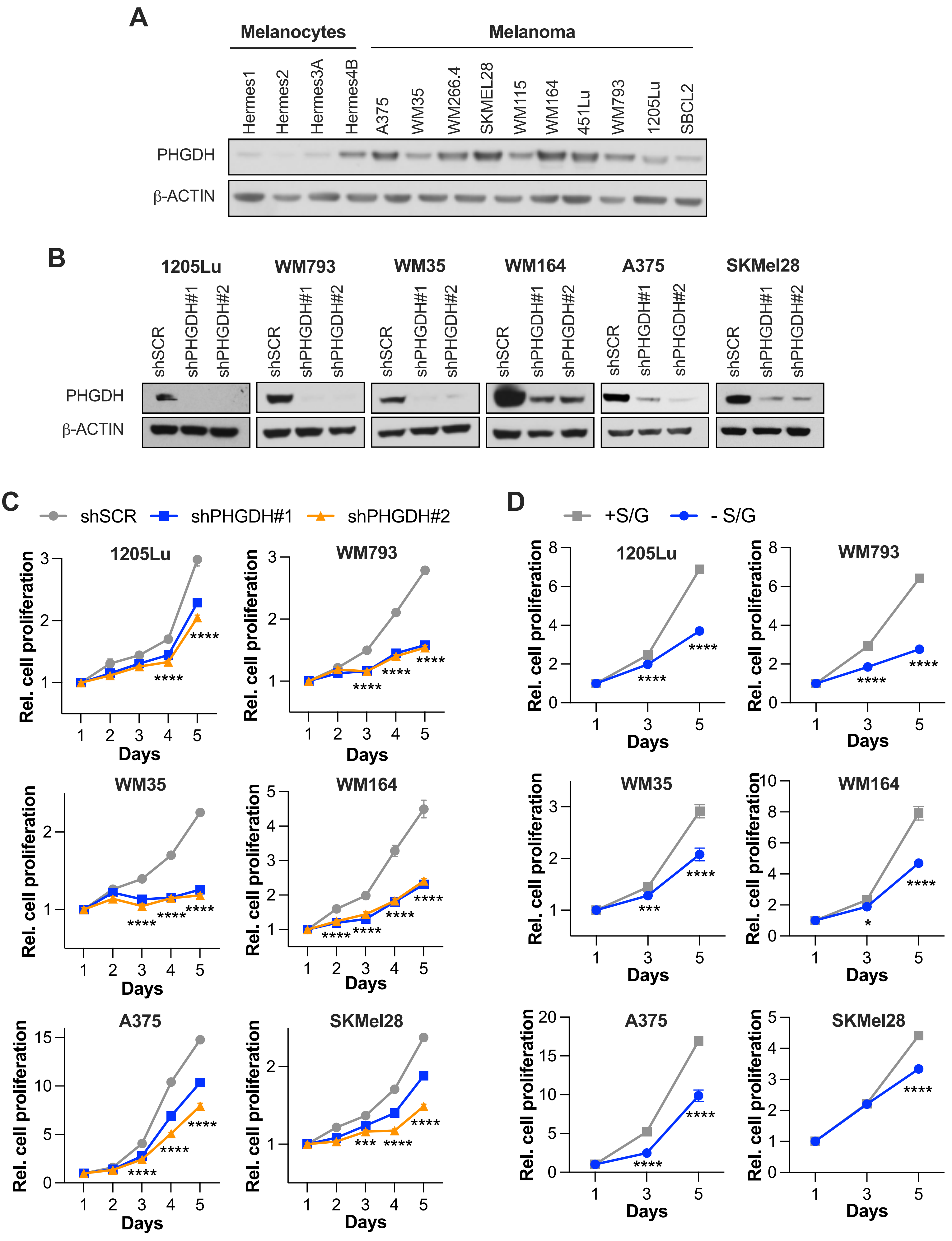


Jasani et al., Supplementary Figure S1

### Supplemental Figure 2

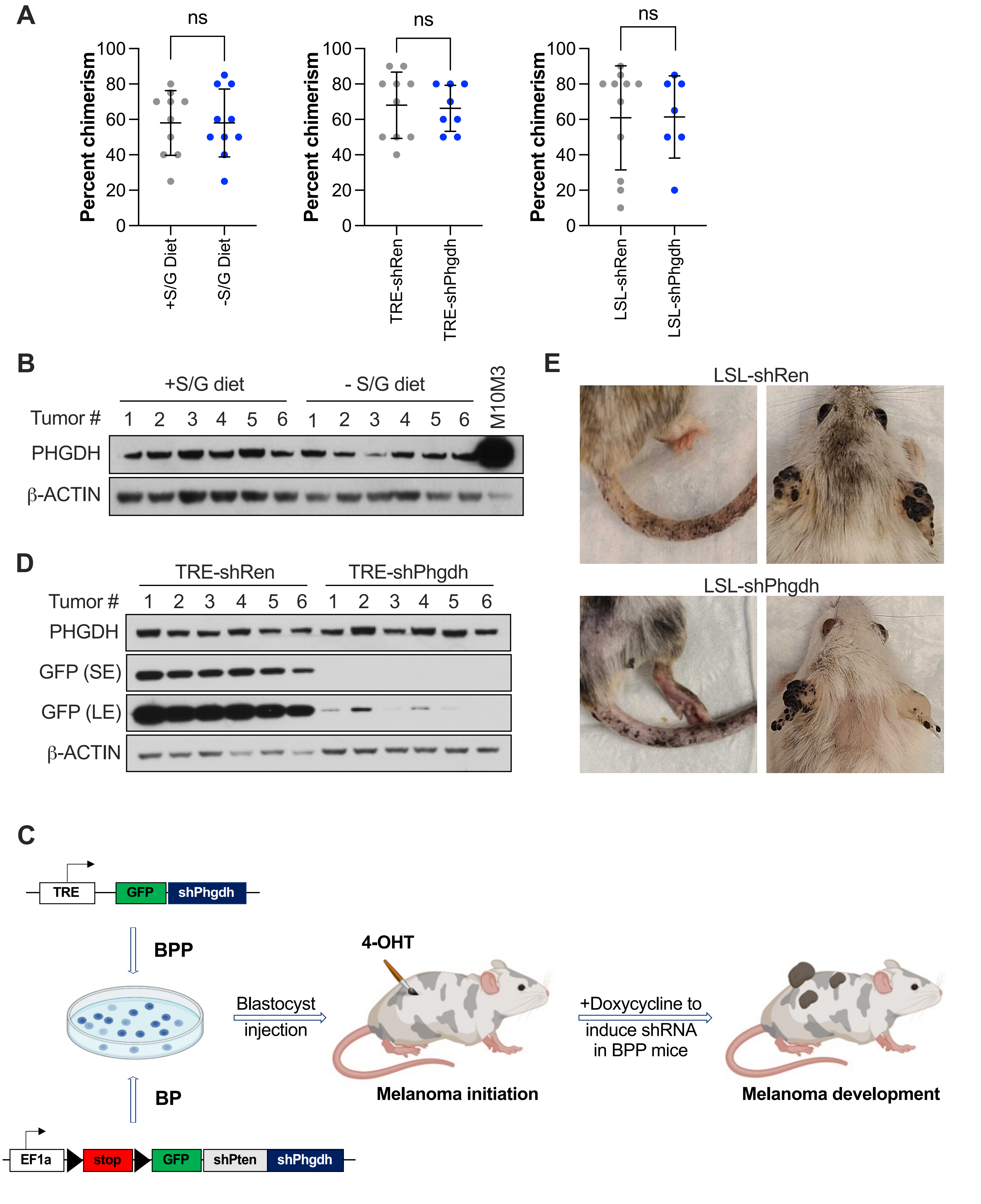


**Jasani et al., Supplementary Figure S2**

### Supplemental Figure 3

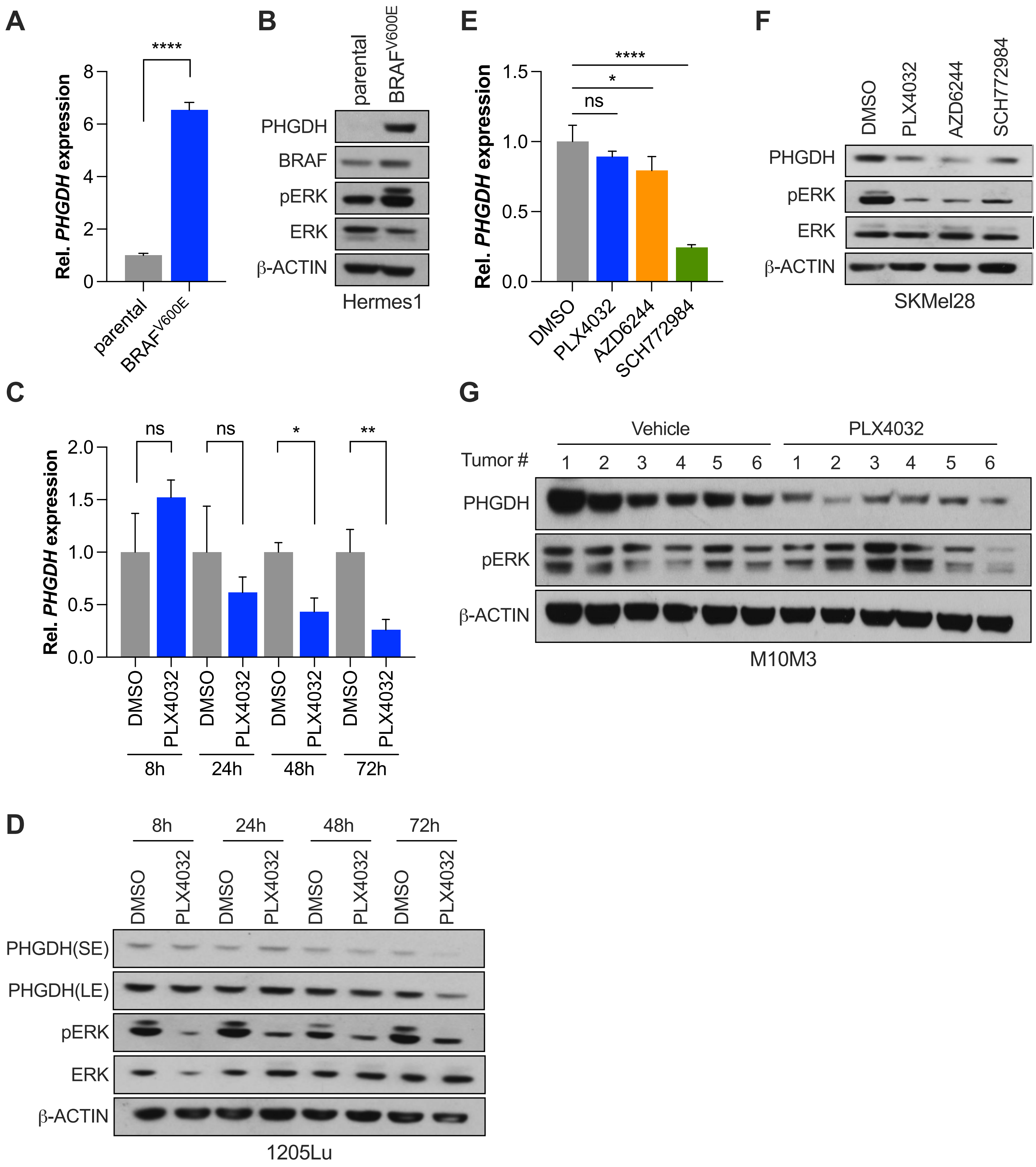


Jasani et al., Supplementary Figure S3

### Supplemental Figure 4

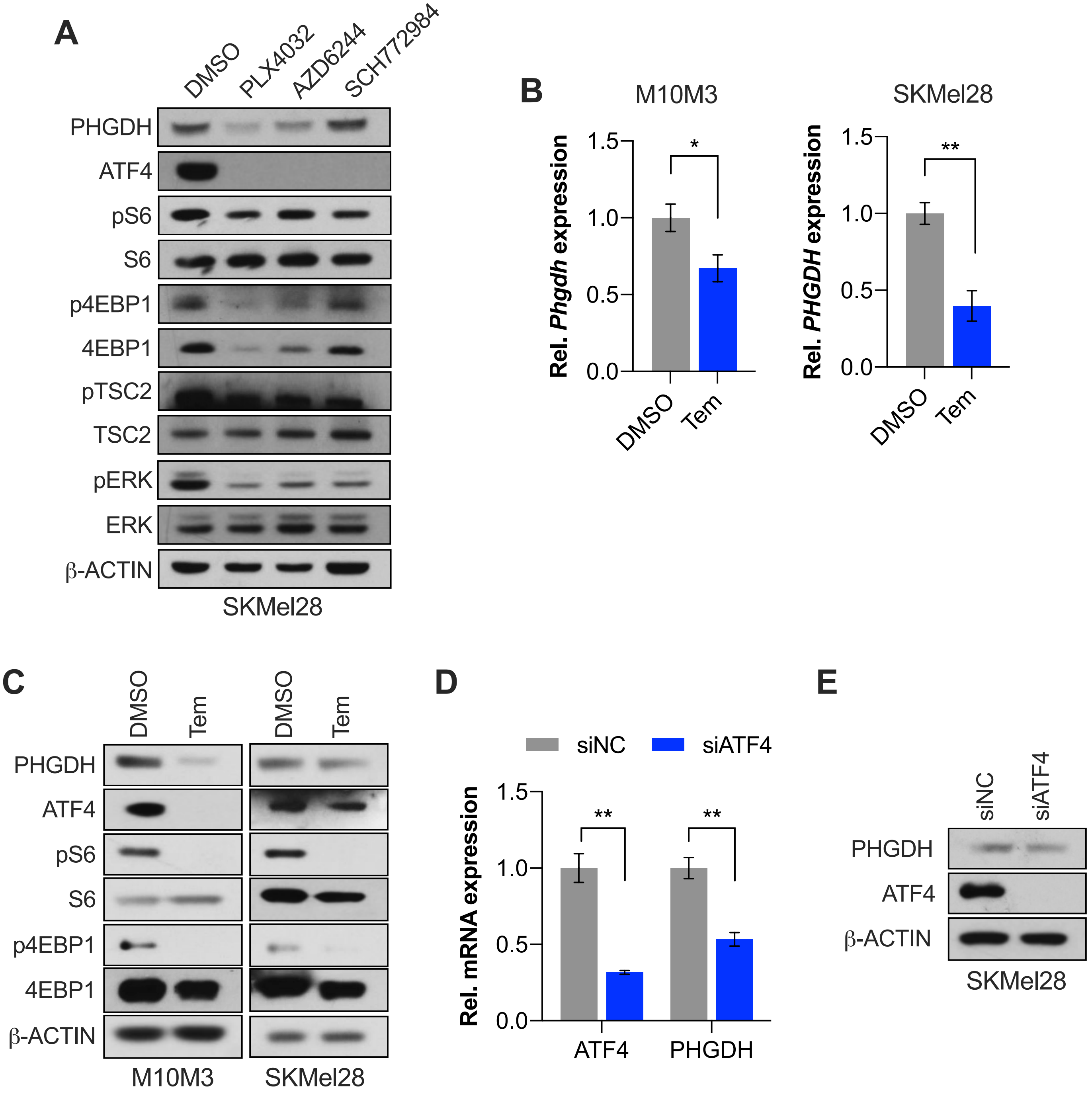


**Jasani et al., Supplementary Figure S4**

### Supplemental Figure 5

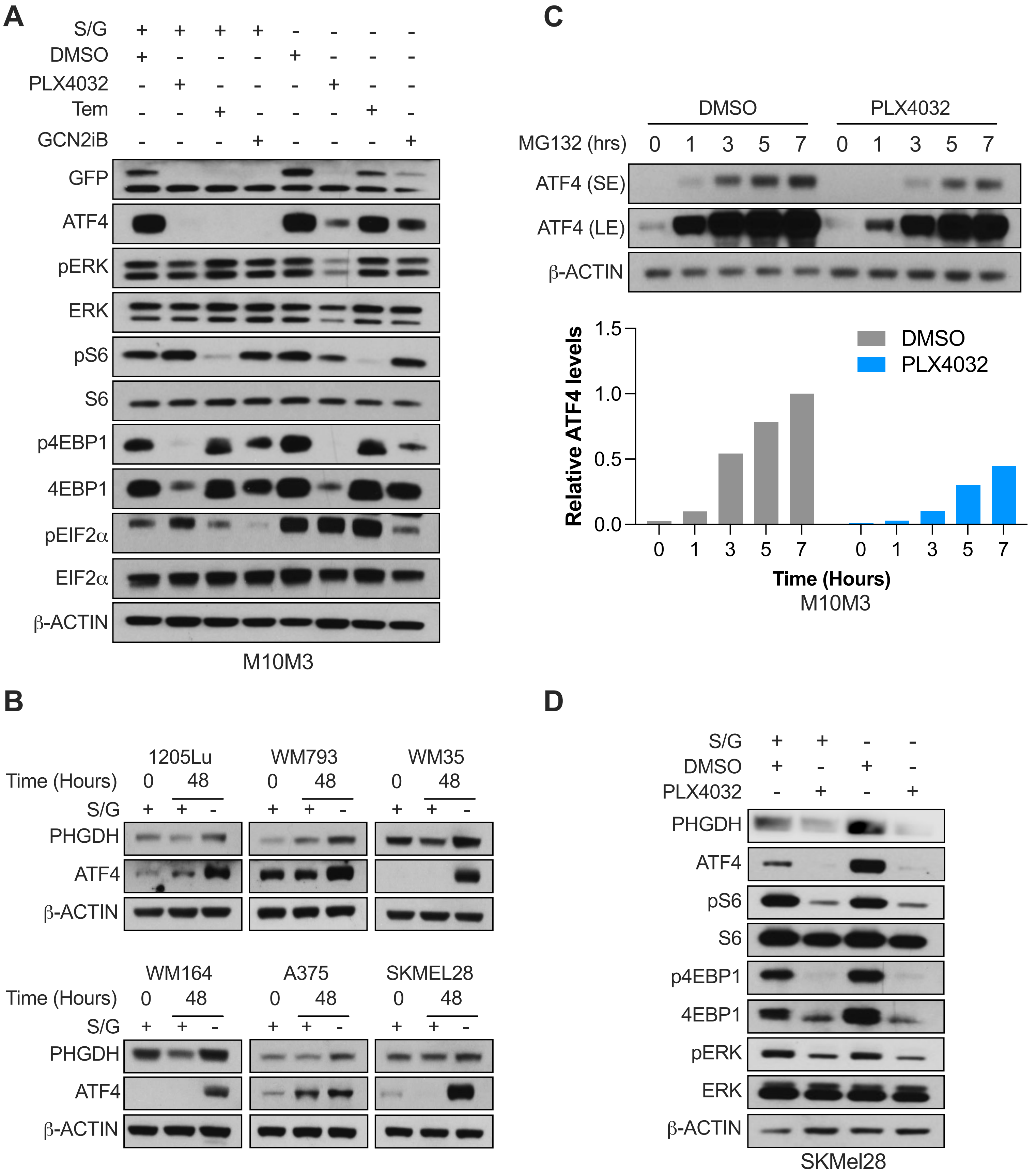


**Jasani et al., Supplementary Figure S5**

### Supplemental Figure 6

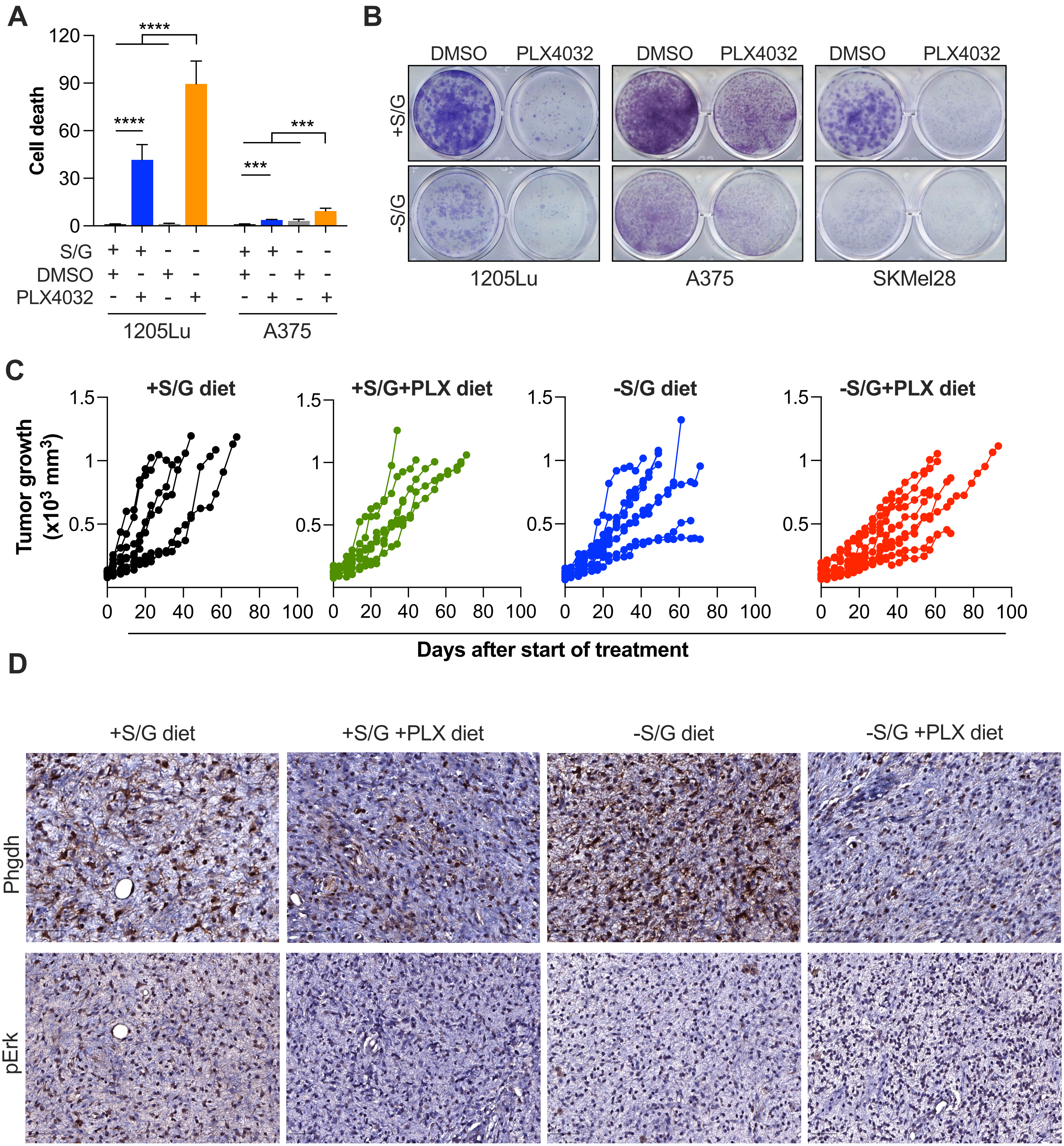


Jasani et al., Supplementary Figure S6
